# Generative cerebral vasculature visualization using spatial transcriptomic data

**DOI:** 10.1101/2025.07.10.664153

**Authors:** Ingrid Berg, Jiqing Wu, Viktor H. Koelzer

## Abstract

The brain is sustained by an intricate vascular network that provides a continuous supply of nutrients and oxygen essential for its function. Understanding the structural variability and region-specific function of this neurovascular system is essential for interpreting neurological alterations in disease models. Leveraging the spatial mRNA-guided generative model Tera-MIND (simulating **Tera**-scale **M**ouse bra**IN**s using a patch-based and boundary-aware **D**iffusion model), we propose to predict the spatial organization of brain vasculature by the co-expression patterns of *Cldn5* and *Acta2* genes and their learned attention map at cell-level resolution. These findings demonstrate the capability of generative AI models trained on single-cell spatial transcriptomic data to reconstruct biologically meaningful higher-order structures. They highlight the potential of such models as in silico systems - GenAI-based simulations that generate realistic representations of biological architecture from spatial molecular data - for high-throughput exploration of vascular function and its dysregulation in neurological disorders. Taken together, our approach repurposes existing spatial transcriptomics datasets to derive new spatial insights into vascular organization, without the need for additional tissue processing.

**Significance Statement:** The brain’s vascular network is complex and functionally essential. Accurate reconstruction of this network at single-cell transcriptomic resolution is key to understanding neurovascular disorders. This study applies spatial transcriptomic and histological data to generate representations of brain vasculature using Tera-MIND, providing a framework that offers a scalable approach for understanding vascular organization.

## Introduction

Neurons rely on continuous blood flow due to their high energy requirements and low capacity for energy storage. Even small disruptions in blood flow can impair neuronal function and survival. Neuronal activity, in turn, modulates blood flow, and several neurological diseases involve vascular dysfunction^1^. Mapping the cerebrovascular network is challenging due to the large variety of blood vessel calibers, from capillaries to large vessels, in a complex 3D structure. However, brain-wide vascular maps are fundamental for understanding how vascular network organization supports neural function and how changes in this system contribute to disease^1^. Recent developments in generative artificial intelligence (GenAI) have supported the in-silico synthesis of realistic biological structures spanning different spatial scales. With complex datasets such as biomedical images and spatial transcriptomics as inputs, GenAI models can learn to reconstruct tissue microanatomy and simulate morphological transitions, establishing *in silico* modeling of whole organisms^2^. In the recent Tera-MIND study, the authors proposed a novel GenAI approach that can capture molecular-to-morphology spatial associations, exemplified by key pathways involved in glutamatergic and dopaminergic neuronal systems^3^. In this application study, we further investigate the potential of generative AI to model biologically meaningful higher-order structures using single-cell spatial transcriptomics data.

## Results

With Tera-MIND, we can visualize an integrated model of gene expression in the neurovascular context. Specifically, we apply the Tera-MIND approach (Fig. 1a and 1b) to generate high-resolution, spatially-resolved visualizations of cerebral microvascular structures, creating a virtual representation of murine brains (exemplified in Fig. 1c and 1d) with projected microvasculature. This enables the spatial analysis of gene expression patterns in relation to vascular architecture and neurovascular function (Fig. 1e-h). In this application study, we focus on two vasculature-associated genes for which spatial transcriptomic information was available in all three mouse brain data sets: *Cldn5* and *Acta2*. Both *Cldn5* and *Acta2* expression play key roles in the function of the neurovascular unit, and dysfunction of these genes is associated with various neurological diseases.

**Figure 1.**
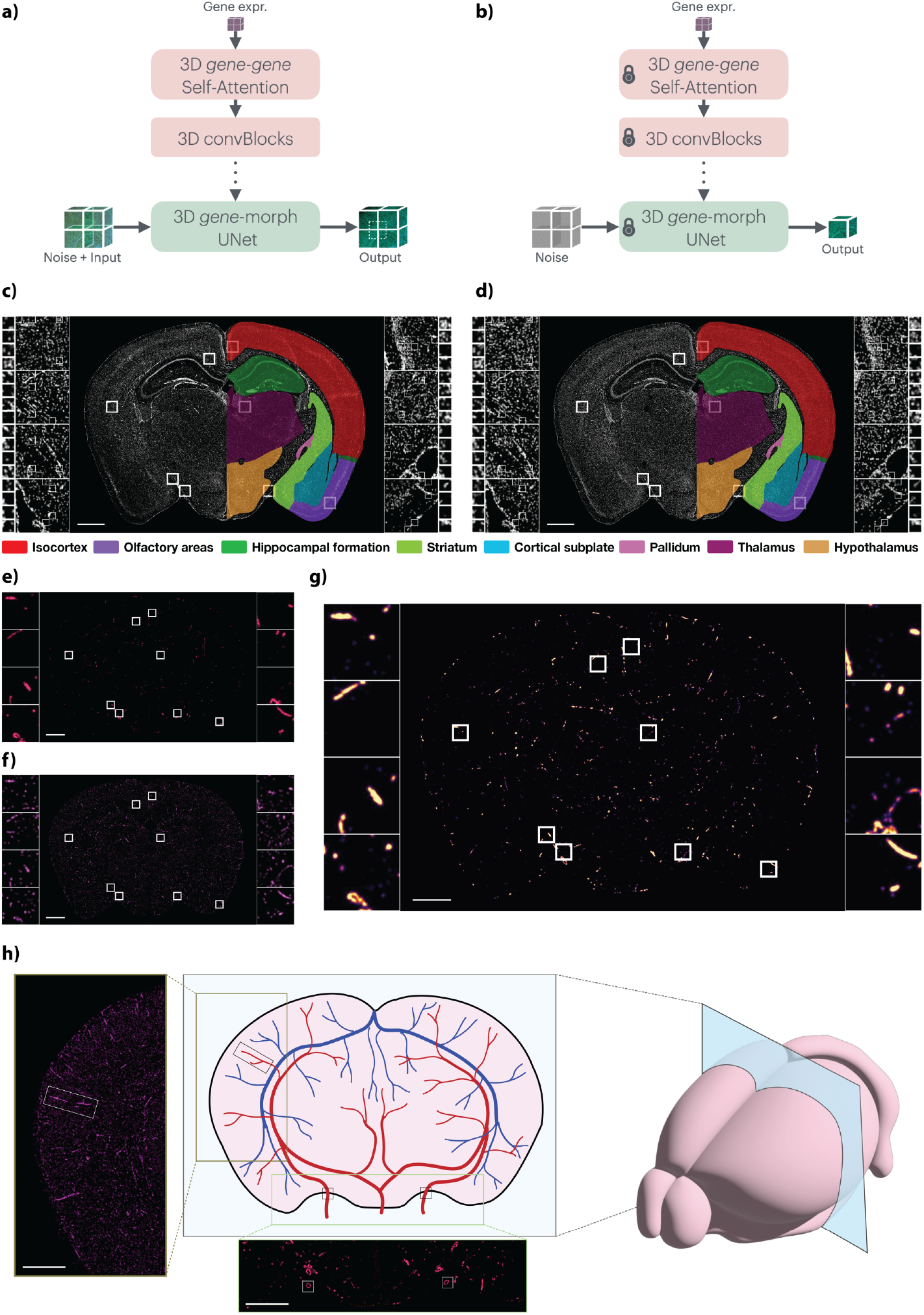
Generative reconstruction of cerebrovascular architecture from spatial transcriptomic data. **a)** Illustration of patch-based diffusion model training. **b)** Illustration of model test/inference. **c)** Image showing the ground truth of a DAPI staining of a coronal section of the mouse brain, including color labels of major brain regions. **d)** Image showing the generated version of the DAPI staining, including color labels of major brain regions. **e)** Generated image showing *Acta2* expression in a coronal brain section. **f)** Generated image showing *Cldn5* expression in a coronal brain section. **g)** Visualization of *Cldn5*-*Acta2* attention. **h)** Schematic illustration of vasculature distribution, including examples of *Cldn5 (left panel)* and *Acta2 (bottom panel)* expression (merged image of 4 consecutive slides) with corresponding structures highlighted with black and white boxes. **c-h)** Length of scale bar (white, bottom left corner) is 1mm.

### 3D modeling of claudin-5 expression in the murine brain

*Cldn5*, a gene encoding the endothelial tight-junction protein claudin-5, is critical for blood-brain barrier (BBB) integrity and is known to exhibit highest expression in brain capillaries and small venules^4^. Using the Tera-MIND model, we predicted *Cldn5* expression across the brain and observed spatial patterns that correspond closely to known vascular anatomy (Fig. 1f). Notably, regions with longitudinally sectioned vessels, such as those extending from the cortex into deeper brain layers^5^, exhibited strong and continuous predicted *Cldn5* signal (Fig. 1h, representative merged image of four adjacent sections), consistent with the expected localization of capillary-rich zones. These findings demonstrate that the model successfully captures biologically meaningful vascular features relevant to BBB-associated gene expression.

### 3D modeling of alpha smooth muscle actin expression

*Acta2*, which encodes alpha smooth muscle actin, is predominantly expressed in vascular smooth muscle cells and marks arterial structures, particularly larger arteries^1^. Tera-MIND predictions of *Acta2* expression localized to regions consistent with major arterial territories, including the ventral surface of the brain where larger arteries are anatomically expected^6^ (Fig. 1e, 1h). To explore spatial interactions between vascular components, we jointly analyzed *Acta2* and *Cldn5* expression using gene-gene attention mapping. This analysis revealed co-localized expression within shared vascular segments (Fig. 1g), suggesting that the model accurately captures complementary features of the vascular hierarchy - from endothelial markers of the blood-brain barrier to smooth muscle-associated arterial identity.

### Vascular network reconstruction

To reconstruct the vascular architecture of the healthy mouse brain, we applied the Tera-MIND model to consecutive 2D spatial transcriptomics sections, focusing on genes associated with the neurovascular unit. The resulting 3D-integrated predictions enabled visualization of vascular continuity and organization across anatomical regions. Fig. 1h illustrates the spatial distribution of vascular features, with predicted *Cldn5* and *Acta2* expression highlighting endothelial and smooth muscle components, respectively. These predictions align with known vascular structures and support downstream analyses of spatial gene expression in the context of vascular integrity.

## Discussion

In this study, we present a generative, spatial gene-informed *in silico* model of the murine cerebral vasculature, reconstructed at teravoxel scale from single-cell spatial transcriptomics data. By leveraging vascular marker genes, the model enables the formation of coherent, spatially-continuous vascular structures that reflect known anatomical and molecular features. This virtual representation supports high-resolution spatial analysis of gene expression in relation to vascular architecture, offering a new framework to study neurovascular organization and function in healthy tissue. Importantly, this work establishes a scalable approach for modeling higher-order tissue structures from single-cell spatial data within their native spatial context, enabling the investigation of gene expression at cellular resolution.

While advanced imaging methods such as light sheet or serial two-photon microscopy provide high-resolution vascular reconstructions, they remain time-consuming and depend on labor-intensive protocols. In contrast, by leveraging the modern high-throughput GenAI approach and reusing spatial transcriptomics datasets, our approach extends their utility by enabling structural analyses, including vascular reconstruction, cell-cell interactions and co-localization. Thus, Tera-MIND provides a complementary avenue for extracting structural insights, extending rather than replacing imaging-based approaches.

Looking ahead, this modeling framework could be extended to incorporate additional modalities and conditions, enabling more comprehensive 3D simulations of tissue organization. Integration with complementary histological, imaging, and genomic datasets may enhance the model’s capacity to capture morphological and functional variability. While this study focuses on healthy brain tissue, the generative approach lays the foundation for future *in silico* exploration of disease-associated perturbations, such as blood-brain barrier dysfunction, once appropriate spatial transcriptomic datasets from disease models become available.

A key strength of our approach lies in its multi-scale ability for interpretable image prediction, enabling detailed visualization of vascular structures from gene expression data. However, a limitation of the current study is that 3D reconstruction is based on ground truth data with noncontiguous 2D sections with 200 micron gaps^7^. This inherent sparsity in the ground truth data introduces challenges for accurate 3D vascular network reconstruction, potentially leading to discontinuities or artifacts. One promising strategy to address this limitation has been shown by Turos *et al*. with X-Pression, which combines single 2D spatial transcriptomics sections with micro-CT imaging to enable full 3D reconstruction of gene expression patterns at the tissue level, eliminating the need for serial sectioning of entire organs^8^.

Guided by the spatial co-expression of vasculature-relevant genes *Cldn5* and *Acta2*, we demonstrated the utility of generative modeling Tera-MIND for vasculature reconstruction using spatial transcriptomics data. This multi-modal approach provides an ethically responsible and cost-effective platform for probing vascular heterogeneity *in silico* and can thereby enable systematic, high-throughput analysis of vascular organization across diverse biological contexts.

## Materials and Methods

### Tera-MIND generative framework

The Tera-MIND model is a novel generative framework designed for tera-scale biomedical data. It leverages paired spatial transcriptomics arrays and co-registered histological images to simulate mouse brains in space. Tera-MIND employs a patch-wise and boundary-aware training strategy, enabling the model to learn from smaller sub-volumes. This design allows for efficient training and inference on high-volume datasets, making whole-brain reconstruction at cell-level resolution computationally feasible under mild hardware requirements^3^. A summary of the generative framework is provided in supplementary methods, with extended methodological details described in Wu *et al*.^3^

### Spatial transcriptomics data

Recent efforts by a network of researchers has resulted in the generation of a detailed cellular atlas of the mouse brain^7,9–11^. The mouse brain data set of coronal sections with high-resolution images of DAPI (4’,6-diamidino-2-phenylindole) and Poly-T (mRNA) stained tissue with the corresponding spatially indexed transcriptomic readout used for development of Tera-MIND was published by Yao *et al*.^7^. Training of Tera-MIND on this data set is described in more detail in the previous study^3^. In short, by training a patch-based and boundary-aware diffusion model on gene expression input and corresponding histological images, it predicts the spatial associations between molecular gene expression markers and morphology.

As described previously^3^, data sets for three different mouse brains were available^7,9^. The primary generation results are based on the data set of a female mouse brain (P56), and the other two available data sets (one female and one male mouse brain, P56) were used as training data. We refer interested readers to the previous study^3^ for a more detailed comparison of ground truth and generated images. Here, we include one example of a ground truth (Fig. 1c) and a corresponding generated (Fig. 1d) DAPI image of a coronal mouse brain section.

### Validation and analysis

After training the model based on the two training data sets with corresponding transcriptomic and histological data, the model of the brain vasculature based on *Acta2* and *Cldn5* expression was generated on the unseen third data set (Fig. 1a and 1b, model training and test). The reconstructed vasculature was validated against available imaging data and known anatomical landmarks from the Xiong *et al*. study^6^.

### Data visualization and figure assembly

Figure illustrations were created and assembled in *Numbers* (version 14.1) and *Adobe Illustrator* (version 29.4).

### Declaration of generative AI and AI-assisted technologies

During the preparation of this work, the authors used ChatGPT (5, OpenAI) to improve the language of the initial manuscript draft. The authors subsequently reviewed and extensively edited the content and take full responsibility for the published article.

## Data Availability

Spatial transcriptomic and histological data (DAPI and Poly-T) for all three mouse brain atlases can be accessed via https://doi.org/10.35077/g.610.

The processed spatial transcriptomic and histological data can be accessed via https://doi.org/10.35077/g.1176.

The code and details on model training and testing are made available at the repository github.com/CTPLab/Tera-MIND.

## Acknowledgments

This study was funded by the Promedica Foundation (F-87701-41-01) and by core funding to the Computational and Translational Pathology Lab led by V.H.K at the University of Basel and University of Zurich.

## Author Contributions

I.B., J.W. and V.H.K. conceived the research idea. J.W. implemented the algorithm and carried out the experiments. I.B., J.W. and V.H.K. analyzed the results. I.B. drafted the manuscript. I.B. and J.W. designed and prepared the figure illustrations. J.W. and V.H.K. critically reviewed and edited the manuscript. J.W. and V.H.K. supervised the project.

## Additional Information

Competing Interest Statement: I.B. and J.W. declare no competing interests. V.H.K. reports being an invited speaker for Sharing Progress in Cancer Care (SPCC) and Indica Labs; advisory board of Takeda; and sponsored research agreements with Roche and IAG, all unrelated to the current study. V.H.K. is a participant of patent applications in computational pathology outside of the submitted work.

## Supplementary Methods

### Tera-MIND generative framework for vascular transcriptomic reconstruction

Tera-MIND is a generative framework that integrates spatial transcriptomics arrays with co-registered histological images to reconstruct brain structures at cellular resolution^1^. In contrast to approaches that detect individual marker genes, Tera-MIND enables the generation of spatial vascular maps by learning molecular-to-morphology associations across entire tissue volumes.

Considering spatial mRNA readouts as three-dimensional “images”, a patch-based diffusion model was applied for efficient training on tera-scale volumetric data, while a boundary-aware denoising component ensured smooth transitions between adjacent patches. Central to the framework is a three-dimensional self-attention block for spatial *gene*-*gene* interactions, which captures correlations among vascular marker genes such as *Cldn5* and *Acta2*. These embeddings are then fed into a UNet-based backbone through cross-attention to guide the reconstruction of vascular morphology.

Training was performed on publicly available mouse brain atlases comprising paired spatial transcriptomics and histological data^2^. Two P56 mouse brains (male and female) were used for model training, and a third unseen P56 female brain served as the held-out test data. This ensured reproducibility and generalization across individuals. Quantitative evaluations against ground truth images were reported in the prior study^1^ and showed high-fidelity reconstructions across multiple scales, with minimal discrepancies measured on cellular features such as nuclear size and cell number^1^.

Despite the tera-scale size of the data (∼0.77 teravoxels), inference is computationally efficient: whole-brain reconstructions can be generated in a patch-wise manner within about a week on a single DGX system equipped with A100 GPUs. Further, the *Cldn5*-*Acta2* guided vasculature mapping takes less than 4 hours.

By seamlessly generating a transcriptomic field rather than classifying isolated spots, Tera-MIND produces coherent vascular hierarchies, from endothelial capillaries to smooth muscle-associated arterial structures. This property enables the visualization of vascular continuity and the analysis of gene-gene interactions within the cerebrovascular network.

For comprehensive architectural and training details, readers are referred to Wu *et al*.^1^.

## References

1. Kirst, C. et al. Mapping the Fine-Scale Organization and Plasticity of the Brain Vasculature. Cell 180, 780-795.e25 (2020).

2. Wu, J. & Koelzer, V. H. Towards generative digital twins in biomedical research. Comput. Struct. Biotechnol. J. 23, 3481–3488 (2024).

3. Wu, J., Berg, I., Li, Y., Konukoglu, E. & Koelzer, V. H. Tera-MIND: Tera-scale mouse brain simulation via spatial mRNA-guided diffusion. Preprint at 10.48550/ARXIV.2503.01220 (2025).

4. Greene, C., Hanley, N. & Campbell, M. Claudin-5: gatekeeper of neurological function. Fluids Barriers CNS 16, 3 (2019).

5. Miyawaki, T. et al. Visualization and molecular characterization of whole-brain vascular networks with capillary resolution. Nat. Commun. 11, 1104 (2020).

6. Xiong, B. et al. Precise Cerebral Vascular Atlas in Stereotaxic Coordinates of Whole Mouse Brain. Front. Neuroanat. 11, 128 (2017).

7. Yao, Z. et al. A high-resolution transcriptomic and spatial atlas of cell types in the whole mouse brain. Nature 624, 317–332 (2023).

8. Túrós, D. et al. Deep learning-based 3D spatial transcriptomics with X-Pression. Preprint at 10.1101/2025.03.21.644627 (2025).

9. Zhang, M. et al. Molecularly defined and spatially resolved cell atlas of the whole mouse brain. Nature 624, 343–354 (2023).

10. Shi, H. et al. Spatial atlas of the mouse central nervous system at molecular resolution. Nature 622, 552–561 (2023).

11. Langlieb, J. et al. The molecular cytoarchitecture of the adult mouse brain. Nature 624, 333–342 (2023).

## References

1. Wu, J., Berg, I., Li, Y., Konukoglu, E. & Koelzer, V. H. Tera-MIND: Tera-scale mouse brain simulation via spatial mRNA-guided diffusion. Preprint at 10.48550/ARXIV.2503.01220 (2025).

2. Yao, Z. et al. A high-resolution transcriptomic and spatial atlas of cell types in the whole mouse brain. Nature 624, 317–332 (2023).

